# Purkinje Cell-specific Deficiency in SEL1L-HRD1 Endoplasmic Reticulum-Associated Degradation Causes Progressive Cerebellar Ataxia in Mice

**DOI:** 10.1101/2024.06.26.600672

**Authors:** Mauricio Torres, Hui Wang, Brent Pederson, Liangguang Leo Lin, Huilun H. Wang, Amara Bugarin-Lapuz, Zhen Zhao, Ling Qi

## Abstract

Recent studies have identified multiple genetic variants of SEL1L-HRD1 ER-associated degradation (ERAD) in humans with neurodevelopmental disorders and locomotor dysfunctions, including ataxia. However, the relevance and importance of SEL1L-HRD1 ERAD in the pathogenesis of ataxia remain unexplored. Here we show that SEL1L deficiency in Purkinje cells leads to early-onset progressive cerebellar ataxia with progressive loss of Purkinje cells with age. Mice with Purkinje cell-specific deletion of SEL1L (*Sel1L^Pcp2Cre^*) exhibit motor dysfunction beginning around 9 weeks of age. Transmission electron microscopy (TEM) analysis reveals dilated ER and fragmented nuclei in Purkinje cells of adult *Sel1L^Pcp2Cre^* mice, indicative of altered ER homeostasis and cell death. Lastly, loss of Purkinje cells is associated with a secondary neurodegeneration of granular cells, as well as robust activation of astrocytes and proliferation of microglia, in the cerebellum of *Sel1L^Pcp2Cre^* mice. These data demonstrate the pathophysiological importance of SEL1L-HRD1 ERAD in Purkinje cells in the pathogenesis of cerebellar ataxia.

**One-sentence summary:** SEL1L-HRD1 ERAD is indispensable for Purkinje cell function and cerebellar ataxia pathogenesis in mice.

## INTRODUCTION

ER-associated degradation (ERAD) is a conserved quality-control pathway responsible for the removal of misfolded proteins in the ER for proteasomal degradation in the cytosol (*1–3*). The suppressor of lin-12-like (SEL1L)-HMG-CoA reductase degradation 1 (HRD1) complex represents the most conserved branch of ERAD from yeast to humans (*4–6*), with SEL1L being the cognate cofactor for the E3 ligase HRD1 (*4, 7, 8*). Using germline- and inducible-knockout (KO) mouse models, we and others have shown that SEL1L-HRD1 ERAD is indispensable for embryonic and adult development *in vivo* (*7, 9–11*). Recent studies using cell type-specific KO mouse models have revealed the vital importance of SEL1L and HRD1 in a cell type- and substrate-specific manner in various physiological processes including food intake, water balance, immune function, stem cell biology, and energy homeostasis (*2, 12–35*). Importantly, we recently identified four pathogenic SEL1L-HRD1 ERAD variants in eleven patients from four families, which are characterized by infantile-onset developmental delay, intellectual disability, microcephaly, facial dysmorphisms, hypotonia, and/or ataxia (*36–38*). We termed this rare neurodevelopmental disorder as ERAD-associated neurodevelopmental disorders with onset in infancy, in short for ENDI (*36*). Moreover, SEL1L S658P variant initially identified in Finnish Hounds with cerebellar ataxia (*39*) was recently reported to cause early onset, non-progressive, ataxia in the mouse model expressing the variant (*40*). While the patients exhibit impaired balance and coordination, the direct role of neuronal SEL1L-HRD1 ERAD in human ataxia remains unknown.

Cerebellar ataxia is a neurological disorder characterized by degeneration of the cerebellum and its associated structures, affecting motor control, balance, and coordination, leading to difficulties in walking, problems with fine motor skills, and speech (*41*). Purkinje cells are a type of neuron found in the cerebellum and are characterized by their elaborate dendritic arbors (*42*). Dysfunction or degeneration of Purkinje cells can lead to a variety of diseases including cerebellar ataxia (*43*). Previous studies in both patients and mouse model systems have implicated several dozen cytosolic and nuclear proteins in the pathogenesis of ataxia, but very few ER proteins (*41, 44*). Mutations in inositol 1,4,5-trispohosphate receptor type 1 (ITPR1), an ER calcium channel. have been identified in early onset, non-progressive, mild spinocerebellar ataxia type 29 (SCA29, OMIM #117360) (*45, 46*). However, the importance of ER protein folding and degradation in the onset and progression of cerebellar ataxia remains unknown.

Here we report an indispensable role of SEL1L, and by extension, SEL1L-HRD1 ERAD, in cerebellar ataxia. Purkinje cell-specific *Sel1L*-deficient mouse model (*Sel1L^Pcp2Cre^*) grow comparably to their wildtype (WT) littermates and appear indistinguishable in gait and balance beam tests at 6 weeks of age; however, they exhibit deteriorating motor function with age, including asymmetric gait, loss of balance, and coordination deficits. Mechanistically, we further show that SEL1L-HRD1 ERAD deficiency leads to a progressive Purkinje cell neurodegeneration with dilated ER and fragmented nuclei. These findings establish SEL1L-HRD1 ERAD as a key player in Purkinje cells in the pathogenesis of cerebellar ataxia.

## RESULTS

### Generation of *Sel1L^Pcp2Cre^* mice with Purkinje cell-specific deletion of SEL1L

To explore the role of SEL1L-HRD1 ERAD in cerebellar ataxia, we generated *Sel1L^Pcp2Cre^* mice with Purkinje cell-specific deletion of Sel1L by crossing *Sel1L^fl/fl^* mice (*7*) with a transgenic mouse expressing Cre recombinase driven by the promoter of Purkinje cell protein 2 (*Pcp2)* (*47*). Pcp2 promoter becomes active starting within the first postnatal week (*48*). Both male and female *Sel1L^Pcp2Cre^*mice grew comparably to *Sel1L^fl/fl^* wildtype (WT) littermates from 3 to 20 weeks of age (Figure 1A-C). Brain weights were comparable between the cohorts at 12-weeks of age but reduced by 10% in *Sel1L^Pcp2Cre^* mice at 20-weeks of age compared to those of WT littermates (Figure 1D-F). Protein levels of both SEL1L and HRD1 were reduced by 60 and 40%, respectively, in total cerebellar lysates from 5-week-old mice (Figure 1G-H). This observation is in line with our previous reports showing the requirement of SEL1L in HRD1 stability (*7*). Deletion of SEL1L in Purkinje cells was further confirmed using confocal microscopy, which showed an 80% reduction of SEL1L signal in Purkinje cells (Figure 1I and J). Protein level of ERAD cofactor OS9 (also an ERAD substrate) was doubled in the cerebellum of 5-week-old *Sel1L^Pcp2Cre^* mice (Figure 1G-H), pointing to SEL1L-HRD1 ERAD dysfunction. By contrast, SEL1L protein level in granule cells was not changed (Figure 1K).

**Figure 1.**
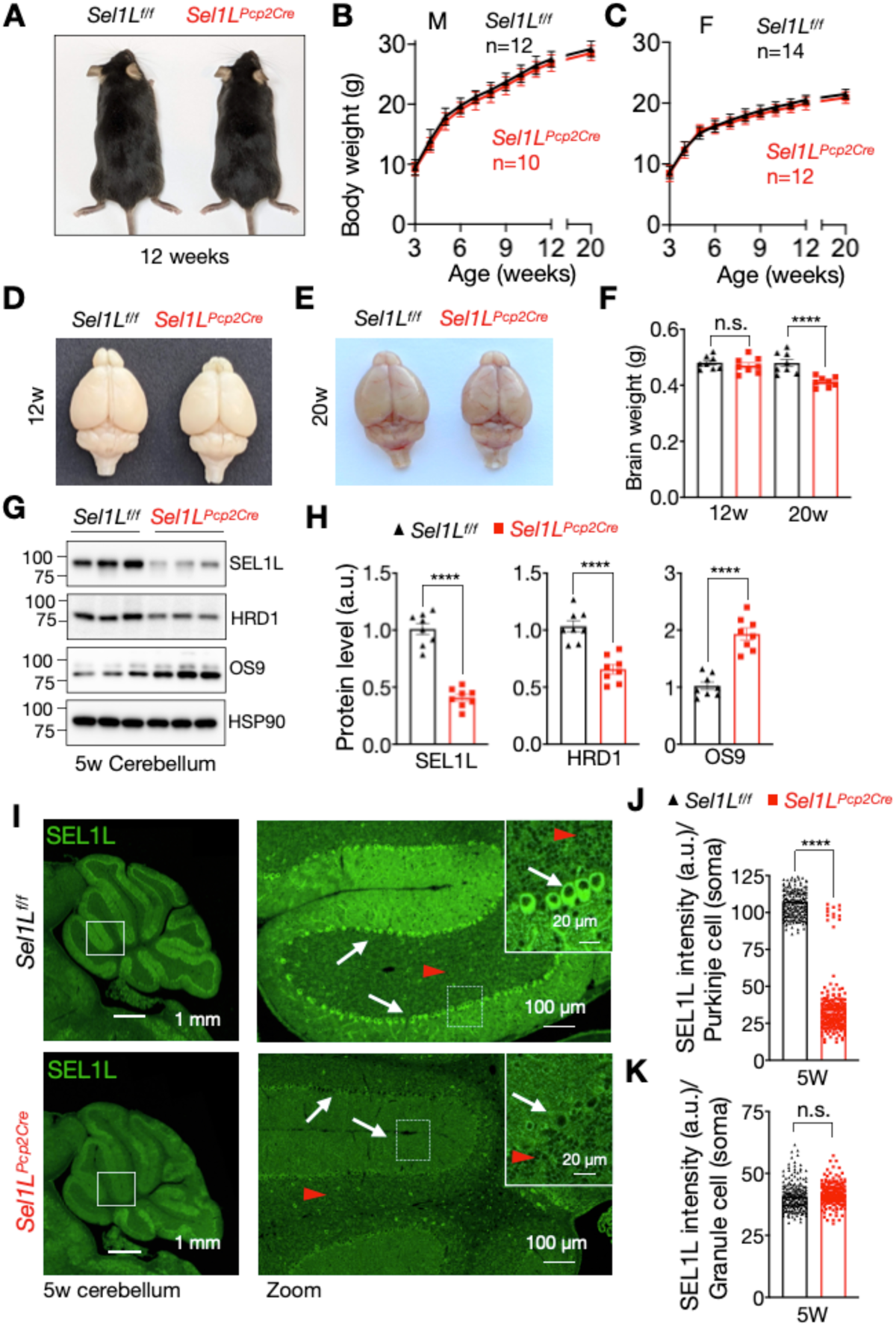
Generation of a mouse model with ERAD deficiency in Purkinje cells. (A) Representative pictures of 12-week-old *Sel1L^f/f^* (Wildtype) and *Sel1L^Pcp2Cre^* (Sel1L knockout) male mice. (B) Growth curve for male and female (C) *Sel1L^f/f^* and *Sel1L^Pcp2Cre^*mice, (3-20 weeks, n = 10-14 mice per group). (D-E) Representative pictures of brains from 12-, and 20-week-old mice. (F) Quantitation of brain weight from 12- and 20-week-old mice (n= 8 mice per group). (G) Western blot analysis of cerebellar protein extracts at 5 weeks of age with quantitation showed in (H) (n= 8 mice per group). (I) Immunofluorescence (IF) of Sel1L protein in cerebellum tissue at 5 weeks of age showing magnifications (insert) for granule cells (red arrow heads) and Purkinje cells (white arrows), with quantitation of Sel1L in the soma of Purkinje cells in (J), and granule cells in (K). Measures obtained from 180-200 cells (n=3 mice per group). Values are mean ± SEM. n.s., not significant; ****p<0.0001 by two-way ANOVA followed by Bonferroni’s multiple comparisons test (B-C), and t-test (F, H, J, and K).

### Purkinje cell SEL1L deficiency leads to progressive cerebellar ataxia

At 6 weeks of age, *Sel1L^Pcp2Cre^* mice displayed similar gait (Figure 2A-C) and performance in balance beam tests (Figure 2D-E and Video 1) as their WT littermates. By 12 weeks, *Sel1L^Pcp2Cre^* mice showed progressively worsening asymmetric gait, evidenced by increased distance between paw match placements and a wider gap between footprints (Figure 2C, video 2, and quantified in Figure 2C). Additionally, 12-week-old *Sel1L^Pcp2Cre^* mice took significantly longer to cross the balance beam (Figure 2D-E and Video 3) and frequently used their tails to prevent falling (Video 3 and Figure 2D). Performance further declined in both tests at 18 and 22 weeks of age (Figure 2C and 2E). Similarly, the hind limb-clasping reflex (*49*) was comparable between the two groups at 6- and 12-weeks but became abnormal at 18 and 22 weeks (Figure 2F-G). Taken together, these data show that Purkinje cell-specific SEL1L-HRD1 ERAD deficiency leads to progressive cerebellar ataxia.

**Figure 2.**
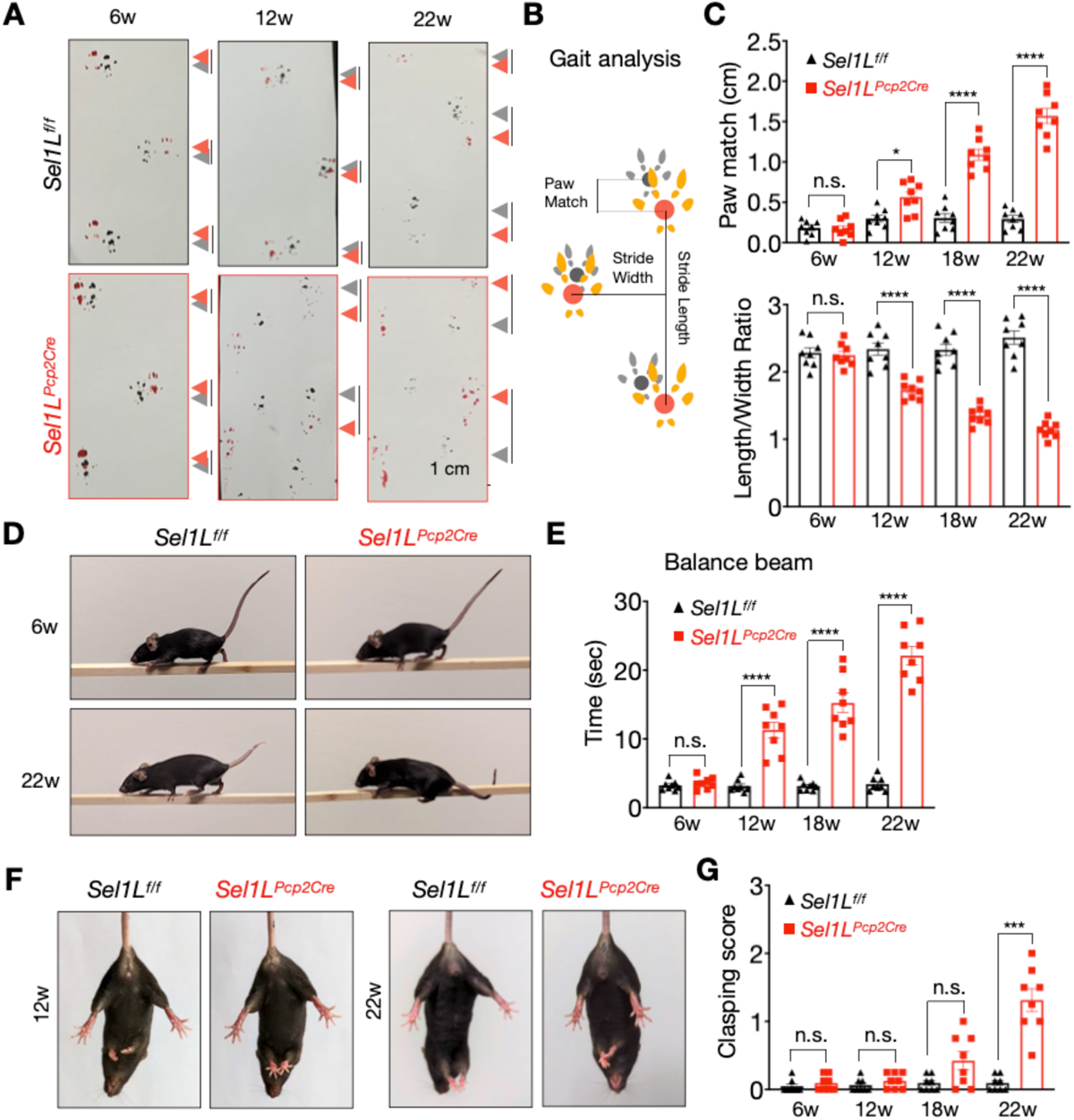
*Sel1L^Pcp2Cre^* KO mice exhibit progressive cerebellar ataxia. (A) Representative pictures of paw prints from 6-, 12-, and 22-week-old mice showing gait analysis for Sel1L^f/f^ and Sel1L^Pcp-2^ mice. The grey and red arrows indicate forelimb and hindlimb, respectively. Lines between grey and red arrow indicate distance between two limbs. (B) Cartoon schematic of the gait analysis. (C) Quantitation of gait analysis of 6 to 22-week-old littermates of both genders (n= 8 mice per group). (D) Representative pictures of balance beam test from 6 and 22-week-old mice. (E) Quantitation of balance beam test from mice at 6, 12,18, and 22 weeks of ages (n= 8 mice per group). (F) Representative pictures of hindlimb clasping of 12- and 22-week-old mice. (G) Quantification of hindlimb clasping from mice at 6, 12,18, and 22 weeks of age (n= 8 mice per group). Values are mean ± SEM. n.s., not significant; ***p<0.001 and ****p<0.0001 by two-way ANOVA followed by Bonferroni’s multiple comparisons test (C, E, and G).

### Progressive loss of Purkinje cells in *Sel1L^Pcp2Cre^* mice with age

Purkinje cells typically exhibit a prominent soma located in a continuous line between the granular and molecular layers of the cerebellum (arrowheads, Figure 3A). Histological examination of the cerebellum revealed comparable Purkinje cell numbers between the cohorts at 5 weeks of age (Figure 3A and 3D), but progressively reduced from 50% to 20% in *Sel1L^Pcp2Cre^* mice at 12- and 20-week of age compared to their WT littermates (Figure 3B-D). Confocal microscopic analyses of the Purkinje cell marker Calbindin further demonstrated the progressive loss of Purkinje cells with age in *Sel1L^Pcp2Cre^* mice (Figure 4A-C), which was supported by Western blot analyses of Calbindin protein levels in the cerebellar lysates (Figure 4E-F). In both assays, Purkinje cell number progressively reduced from 40-50% at 12 weeks, to 10-20% at 20 weeks in *Sel1L^Pcp2Cre^* mice, compared to those of WT littermates (Figure 4D and 4F). Concomitantly, we observed a nearly 40% reduction in the number of granule cells (Figure 3C and E), and a 60% reduction in the molecular layer of *Sel1L^Pcp2Cre^*mice at 20 weeks of age (Figure 3F-G), in line with the known effects of Purkinje cell deficiency on granule cells and molecular layer (*50–52*).

**Figure 3.**
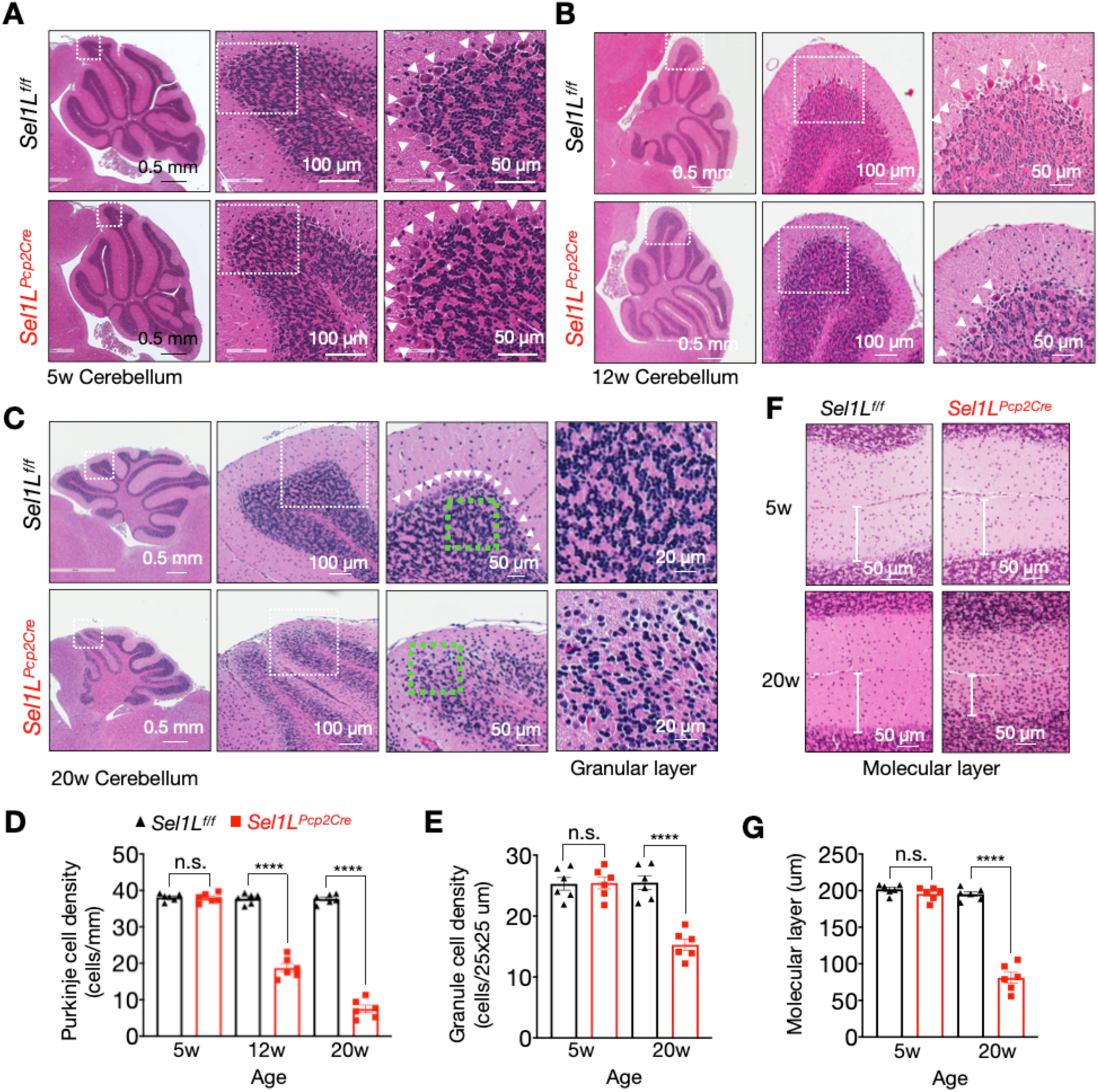
Purkinje cell *SEL1L* deletion leads to Purkinje cell loss and degeneration of the granular layer. (A-C) Hematoxylin and eosin (H&E) stained sagittal sections of the cerebellum from 5-(A), 12-(B), and 20-week-old mice (C). White arrowheads indicate Purkinje cells. (D) Quantitation of Purkinje cell density in H&E-stained sagittal sections of the cerebellum of mice at 5, 12, and 20 weeks of age (n= 6 mice per group). (E) Quantitation of granular cells in an area of 25 µm x 25 µm on the granular layer of the cerebellum (n= 6 mice per group). (F) Representative images of the molecular layer (ML) at 5- and 20 weeks of age, with quantitation of ML thickness shown in G (n= 6 mice per group). Values are mean ± SEM. n.s., not significant; ****p<0.0001 by two-way ANOVA followed by Bonferroni’s multiple comparisons test (D, E and G).

**Figure 4.**
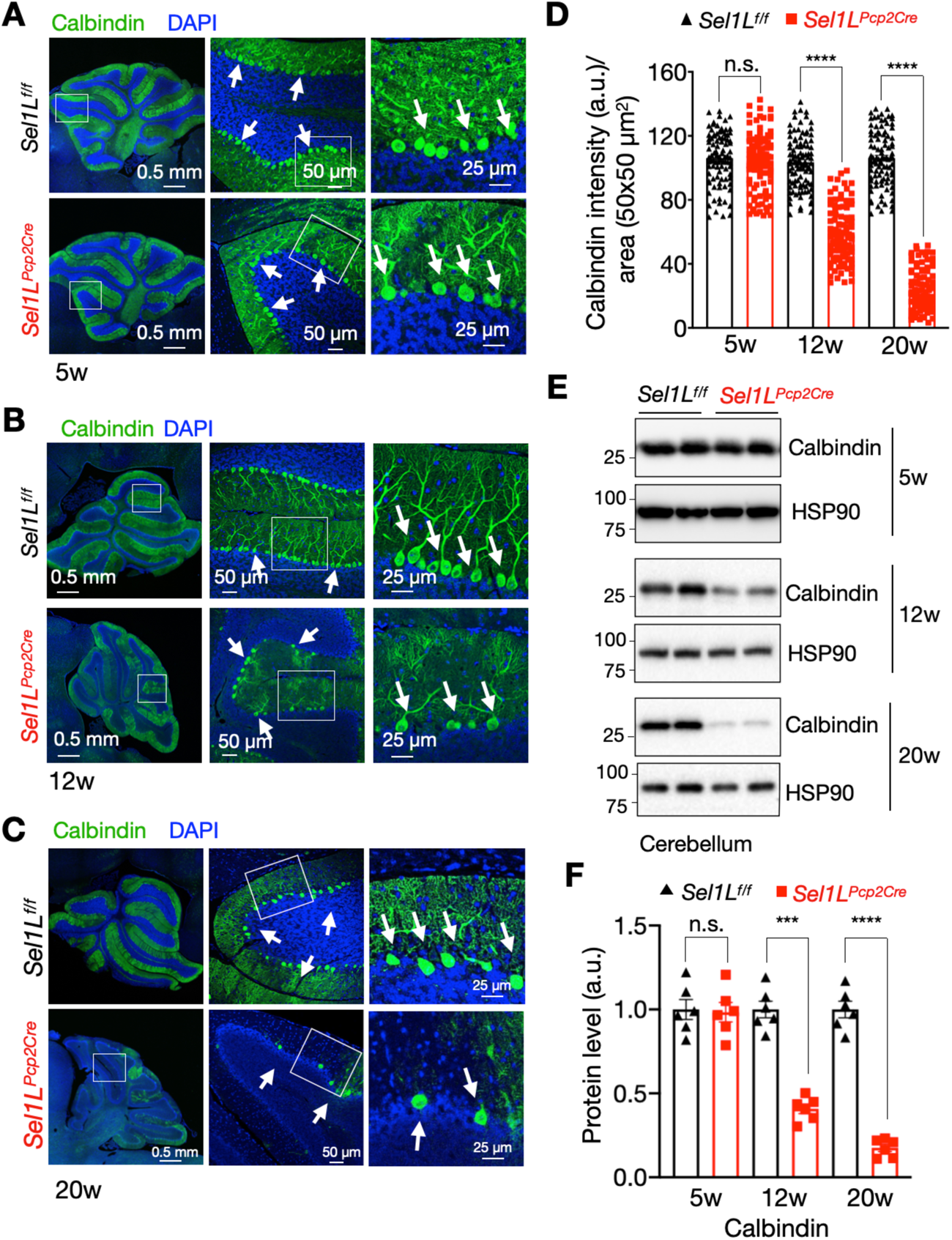
SEL1L deletion leads to a progressive reduction of calbindin-positive Purkinje cells in the cerebellum. (A-C) Representative confocal images of Calbindin (green) and DAPI (blue) staining in the cerebellum of 5-(A),12-(B), and 20-(C) week-old mice. White arrows indicate calbindin-positive Purkinje cells. The magnification of the selected regions is showed in the lateral panel for each figure. (D) Quantitation of calbindin signal intensity in the cerebellar cortex of 5-,12-, and 20-week-old mice (total of 180-200 cells from n=3 mice each cohort). (E) Western blot analysis of Calbindin and HSP90 proteins in protein extracts from cerebellum of 5-, 12- and 20-week-old mice. Quantitation of calbindin levels normalized to loading control is showed in (F) (n= 6 mice per group). Values are mean ± SEM. n.s., not significant; *p<0.05, **p<0.01, ***p<0.001 and ****p<0.0001 by two-way ANOVA followed by Bonferroni’s multiple comparisons test (D and F).

### ER expansion and Purkinje cell neurodegeneration in the absence of SEL1L

We next evaluated the ER ultrastructure of Purkinje cells in the cerebellum of 5- and 9-week-old mice using transmission electron microscopy (TEM). At 5 weeks of age, overall ultrastructural of Purkinje cells of *Sel1L^Pcp2Cre^*mice were largely normal, including the ER, albeit being slightly dilated, compared to that of WT littermates (Figure 5A, quantitated in Figure 5D). At 9 weeks of age, dramatic changes have taken place in Purkinje cells of *Sel1L^Pcp2Cre^*mice: dilated ER (quantified in Figure 5D), fragmented nuclei and the presence of electrodense particles in the cytosol, possibly associated with the accumulation of protein aggregates (Figure 5B-C). In addition, terminal deoxynucleotide transferase-mediated dUTP nick end labeling (TUNEL) assay showed elevated cell death in both Purkinje cell and granular layers of *Sel1L^Pcp2^* mouse at 12 weeks of age (Figure 5E-F), but not at 5 weeks of age (data not shown). These results show that SEL1L deficiency in Purkinje cells leads to a progressive organellar dysfunction and cell death with age.

**Figure 5.**
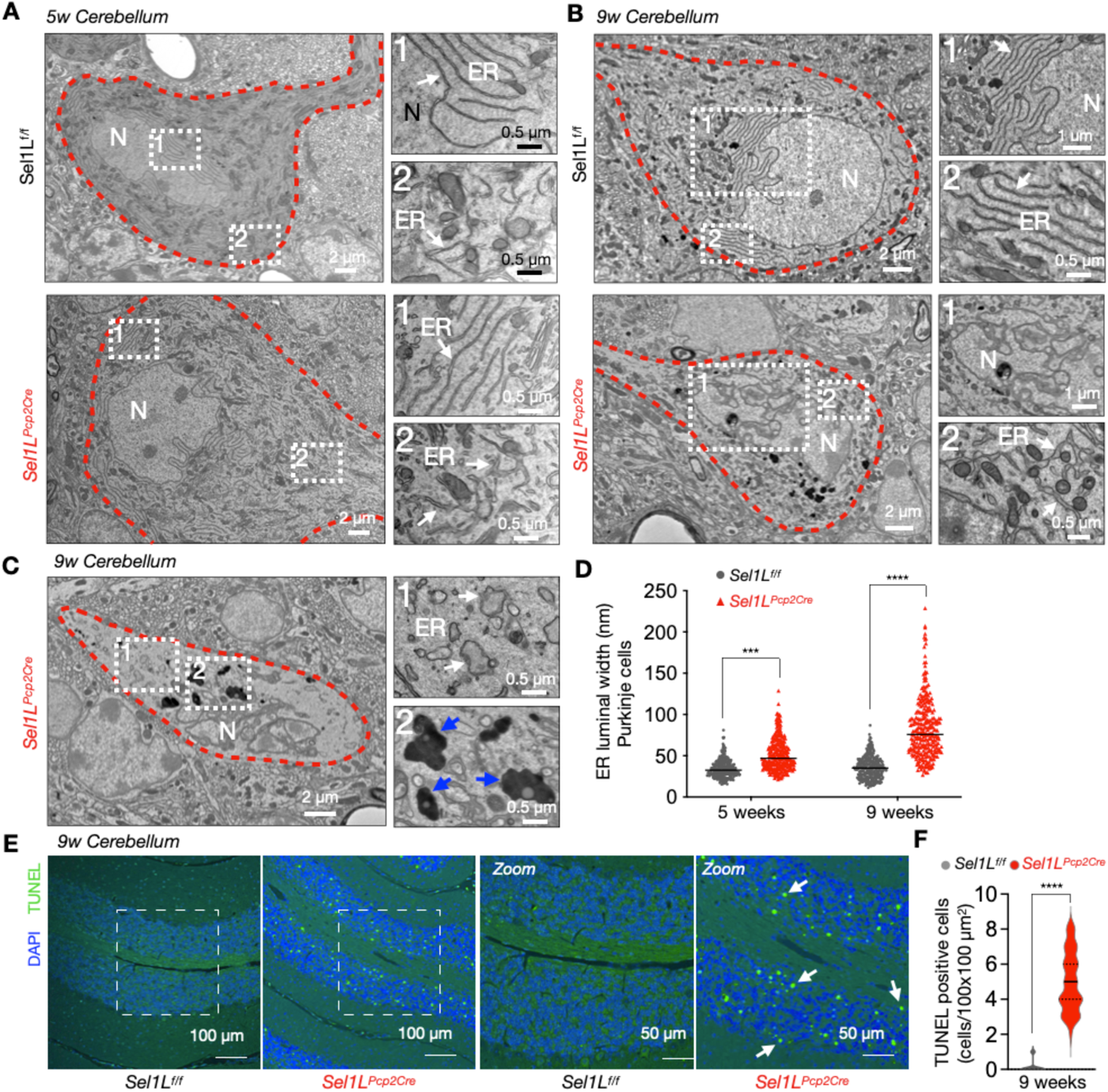
ER expansion and Purkinje cell neurodegeneration in *Sel1L^Pcp2Cre^* mice. (A) Representative TEM images of wild type and Sel1L-deficient Purkinje cells (highlighted by red dotted lines) from 5-week-old mice. Inserted panel 1 and 2 are showing magnification of the endoplasmic reticulum (ER, white arrows) in two different locations of the cell (n= 3 mice per group). (B) Representative TEM images of wild type and Sel1L-deficient Purkinje cells showing fragmentation of the nucleus (N), and ER expansion at 9 weeks of age (n= 3 mice per group). (C) Representative TEM images of Sel1L-deficient Purkinje cells showing ER expansion and accumulation of electrodense structures (blue arrows) at 9 weeks of age (n= 3 mice per group). (D) Quantification of luminal ER width in TEM images of mice at 5 and 9 weeks of age (250-300 measurements from 10 cells, n=3 mice per group). (F) Representative fluorescence images of TUNEL-labeled and DAPI-stained sagittal cerebellar sections of *Sel1L^f/f^* and *Sel1L^Pcp2^* mice at 9 weeks of age. Magnification of selected areas are showed in the left panels. White arrows indicate TUNEL positive labeled nuclei. (F) Quantitation of TUNEL positive cells in the cerebellum of 9-week-old mice (n=3 mice per group). Values are mean ± SEM. n.s., not significant; ***p<0.001 and ****p<0.0001, by two-way ANOVA followed by Bonferroni’s multiple comparisons test (D), and t-test (F).

### Elevated astrocyte activation and presence of microglia in *Sel1L*-deficient cerebellum

During the neurodegenerative process, damaged neurons release inflammatory signals that can activate glial populations, such as GFAP-(glial fibrillary acidic protein)-positive astrocytes (known as Bergmann glia in the cerebellum), and IBA1-(Ionized calcium-binding adaptor molecule 1) positive microglial cells (*53, 54*). Confocal microscopic analyses revealed a marked increase in GFAP signals in areas with loss of Purkinje cells in 20-week-old *Sel1L^Pcp2^* mice (Figure 6A). This finding was further supported by Western blot analysis, which showed a 4-fold increase of GFAP protein levels in *Sel1L^Pcp2^* mice at 12 and 20 weeks of age, but not at 5 weeks of age, compared to that in WT littermates (Figure 6B-C). Moreover, the number of microglial cells was progressively increased from 5- to 12-week-old *Sel1L^Pcp2^*mice, accompanied by hypertrophy of cell bodies and thickening of their processes (Figure 6D-F). Taken together, these data demonstrate that SEL1L deficiency in Purkinje cells leads to not only the loss of Purkinje cells, but also a concomitant increase of neuroinflammation within the cerebellum.

**Figure 6.**
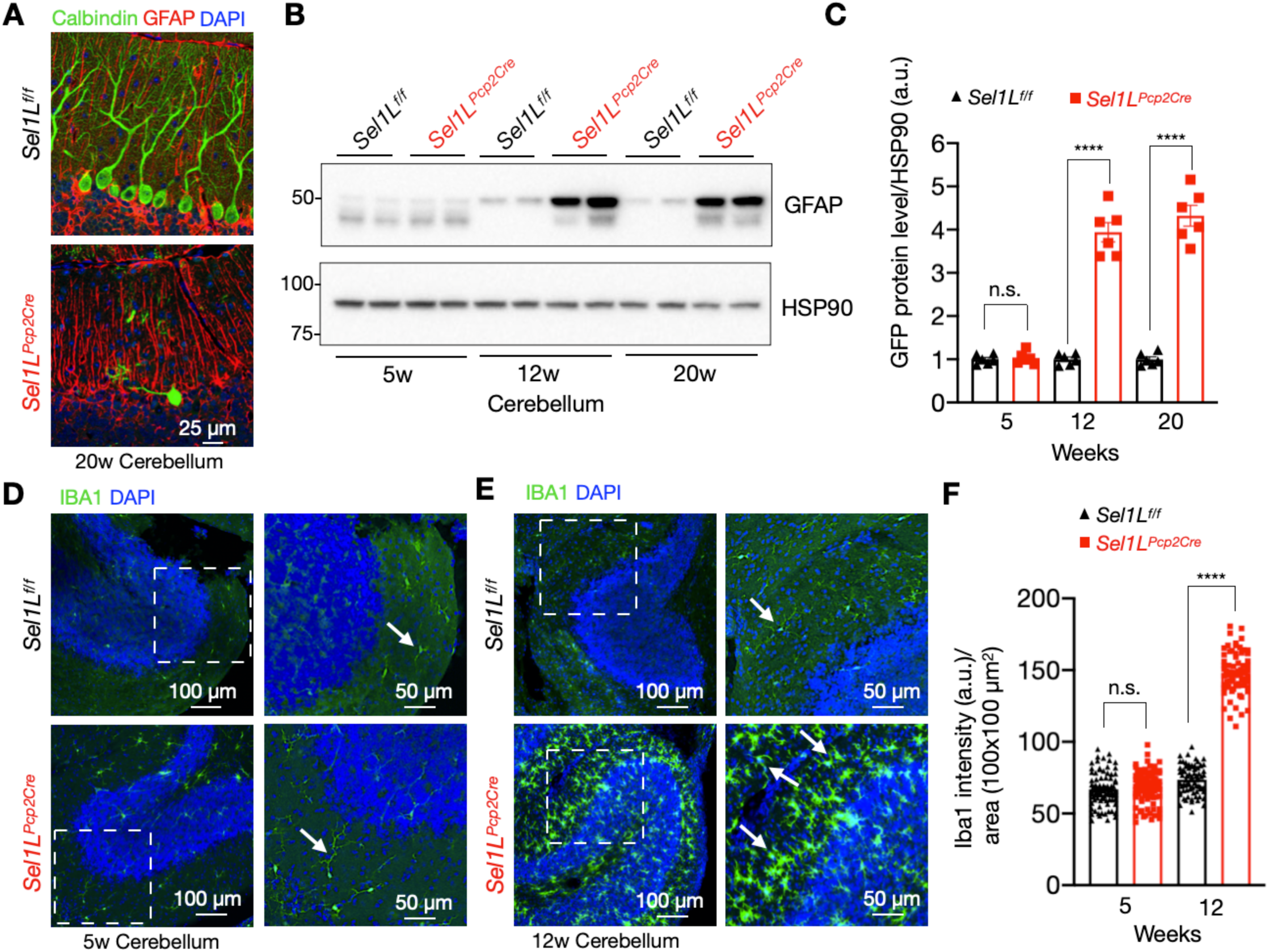
Elevated astrocyte activation and microglia proliferation in *Sel1L^Pcp2Cre^* mice. (A) Confocal images of Purkinje cells stained with Calbindin (green) and GFAP for astrocyte cells (red) in *Sel1L^f/f^* and *Sel1L^Pcp2^* mice at 20 weeks of age (n= 3 mice per group). (B) Western blot analysis of GFAP in the cerebellum of 5, 12, and 20-week-old mice with quantitation shown in (C), (n= 6 per group). (D) Confocal images of IBA1-positive microglia (green) and DAPI (blue) staining in the cerebellum of 5-(D), and 12-(E) week-old mice (n= 3 mice per group). (F) Quantitation of IBA1 positive microglia cell in the cerebellum of 5-,12-, and 20-week-old mice. Values are mean ± SEM. n.s., not significant; *p<0.05, **p<0.01, ***p<0.001 and ****p<0.0001, by two-way ANOVA followed by Bonferroni’s multiple comparisons test (C and F).

### Altered ER homeostasis in *Sel1L*-deficient Purkinje cells

Lastly, we examined the ER homeostasis in the *Sel1L^Pcp2^*mice at 5 weeks of age. The unfolded protein response (UPR) and ERAD are two essential components of the protein quality-control system (*2*). The accumulation of unfolded or misfolded proteins in the ER, induces the activation of URP sensors such as IRE1⍺ and PERK. While IRE1⍺ protein level was moderately elevated (Figure 7A), there was no significant increase in IRE1α-mediated splicing of the downstream effector *Xbp1* mRNA in the cerebellum of *Sel1L^Pcp2^*mice at 5 weeks of age (Figure 7C-D). Similarly, while PERK protein level was significantly increased in the cerebellum of *Sel1L^Pcp2^* mice (Figure 7A-B), there was no significant changes in the expression of its downstream effectors such as phosphorylation of eIF2⍺, ATF4, and CHOP (Figure 7A-B). Additionally, protein levels of the ER chaperones BiP and GRP94 were significantly elevated in the cerebellum of *Sel1L^Pcp2^* mice at 5 weeks of age (Figure 7E-F). This finding was further corroborated by confocal microscopic analyses showing increased KDEL (ER chaperones) and BiP protein levels in Purkinje cells of *Sel1L^Pcp2^* mice (Figure 7G-J). Hence, we conclude that SEL1L deficiency resets ER homeostasis in Purkinje cells.

**Figure 7.**
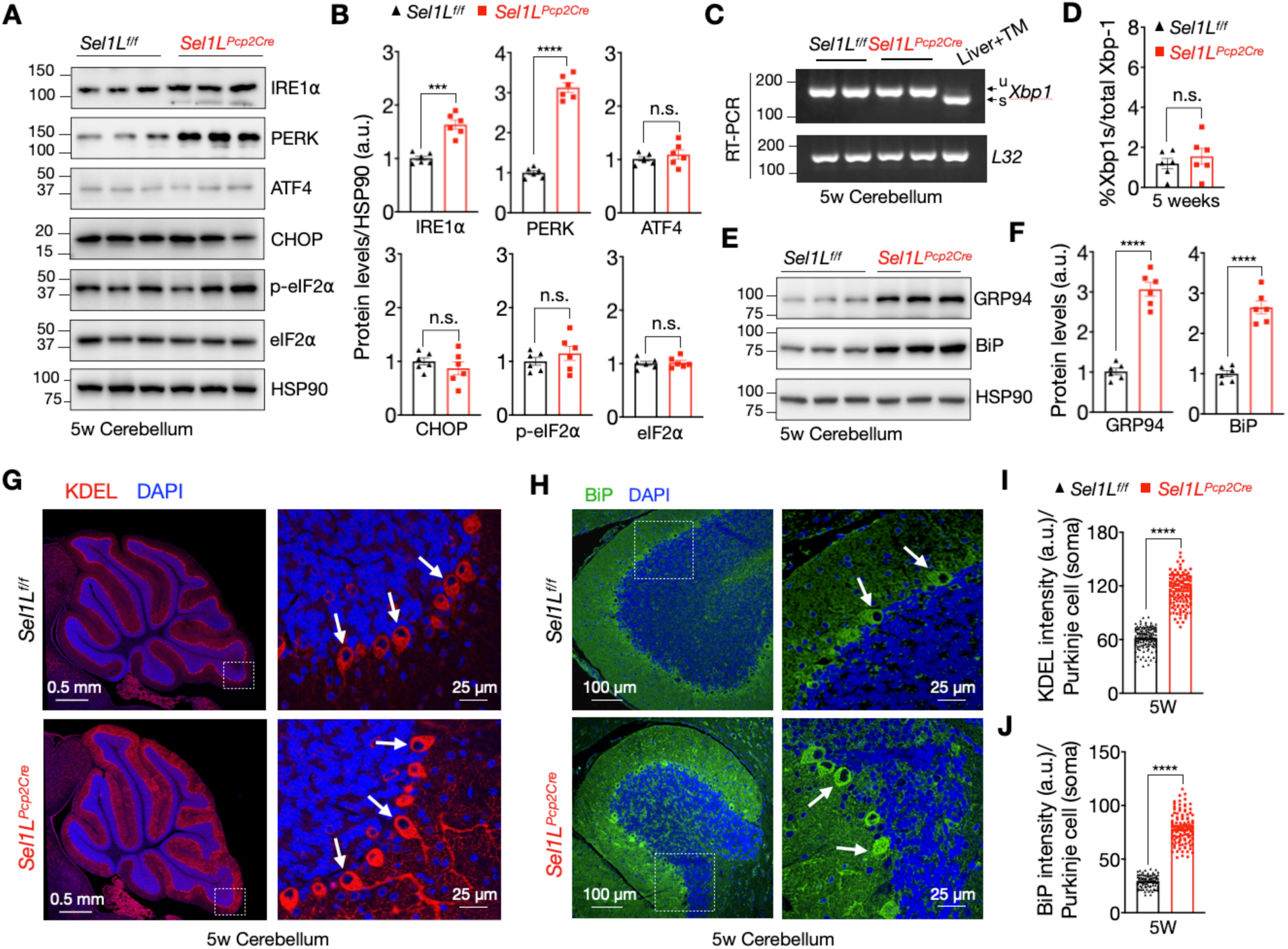
Sel1L deficiency is associated with mild UPR in Purkinje cells of *Sel1L^Pcp2Cre^* mice. (A) Western blot analysis of the UPR sensors IRE1⍺ and PERK, including ATF4, CHOP, and eIF2⍺ pathways in the cerebellum of mice at 5 weeks of age, with quantitation is shown in (B), (n=6 mice per group). (C) RT-PCR of *Xbp1* mRNA splicing in the cerebellum of 5-week-old mice. u and s indicate the unspliced and spliced form of *Xbp1*. Liver treated with tunicamycin (TM, ER stress inducer) is used as a positive control. Quantitation of the spliced form of Xbp1 (Xbp1s) is shown in (D), (n=6 mice per group). (E) Western blot analysis of the ER-chaperones GRP94 and BiP in cerebellum from 5-week-old mice with quantitation shown in (F), (n= 6 mice per group). (G) Representative confocal images of KDEL (red signal), and BiP (green signal, shown in H), in the cerebellum of 5-week-old mice. White arrows indicate Purkinje cells (n= 3 per group). (I) Quantitation of KDEL and BiP signal intensity (J) in the soma of Purkinje cells (total of 120-150 cells from n=3 mice each cohort). Values are mean ± SEM. n.s., not significant; *p<0.05, **p<0.01, ***p<0.001 and ****p<0.0001, by t-test.

## DISCUSSION

Our recent studies identified several patients carrying the SEL1L or HRD1 variants with hypotonia or ataxia (*36, 37*), but the role of SEL1L-HRD1 ERAD in the pathogenesis of cerebellar ataxia remain unknown. Using a Purkinje cell-specific KO mouse model, here we provide definitive evidence for the essential role of SEL1L-HRD1 ERAD in Purkinje cell neurodegeneration and cerebellar ataxia in mice. *Sel1L^Pcp2^* mice develops early-onset, progressive cerebellar ataxia characterized by a progressive loss of Purkinje cells, neuroinflammation, and cerebellar degeneration – mirroring the characteristics of cerebellar ataxia in patients (*43*). This study demonstrates that SEL1L-HRD1 ERAD in Purkinje cells play an essential role in the pathogenesis of cerebellar ataxia.

Purkinje cells are some of the largest neurons in the brain with a prominent ER structure that form an interconnected network regulating calcium signaling, protein synthesis, folding, and protein trafficking (*55*). Previous studies have shown that Purkinje cells are particularly vulnerable to alterations in ER homeostasis due to their high demand for protein synthesis needed to maintain a large number of synapsis (*56, 57*). Our data indicate that Purkinje cells cannot adapt to a significant reduction in SEL1L-HRD1 ERAD function, ultimately leading to cell death. One potential detrimental effect of SEL1L-HRD1 ERAD loss in Purkinje cells could be the accumulation of misfolded proteins in the ER, resulting in cell death due to chronic ER stress. In this context, we observed an accumulation of ER chaperones such as BiP and Grp94, and evidence of dilated ER by TEM. However, we did not detect an active UPR, as indicated by the absence of Xbp-1 mRNA splicing, a downstream effector of IRE1⍺, or changes in ATF4, CHOP or eIF2⍺, three common downstream effectors of the PERK pathway. These results may suggest that additional mechanism independent of ER-stress may contribute to the degeneration and cell death of Purkinje cells. In previous studies, we reported pathological changes associated with the accumulation and ER retention of ERAD substrates (*12, 15, 20, 26, 32, 33, 58–60*), and mitochondria dysfunction in ERAD-deficient cells (*30*). We hypothesize that alterations in the biosynthesis of specific factors involved in maintaining active synapsis, cell-cell interactions, or the reduction of active neurotrophic receptors could also contribute to Purkinje cell death (*61, 62*). We are currently exploring different approaches to isolate and identify ERAD-specific substrates in Purkinje cells. Future studies using single-cell analysis could help identify specific alterations of Purkinje cells, granular cells and glial cells during the progression of cerebellar ataxia in *Sel1L^Pcp2^* mice.

We also observed compelling evidence of robust astrocyte activation and an elevated number of microglia associated with the loss of Purkinje cells in the cerebellum of Sel1L^Pcp2Cre^ mice. Bergmann glia, an unusual type of astrocyte, are closely associated with Purkinje cells, enclosing both soma and synapses and establishing complex anatomical and functional interactions (*63*). The typical radial appearance of Bergmann glia and normal dendritic tree pattern of Purkinje cells was observed in 5-week-old animals, suggesting a normal postnatal development of the cerebellum. Normal levels of GFAP and IBA1 markers were detected in 5-week-old mice, suggesting absence of neuroinflammation at this age. However, at 12 weeks of age, we found elevated levels of GFAP and an increased number of microglia in Sel1L^Pcp2Cre^ mice, which correlated with a reduction in the number of Purkinje cells and the development of ataxia. We suspect that dysfunctional Purkinje cells could release inflammatory signals that trigger the activation of Bergmann glia. Activated astrocytes may lead to alterations in neurotransmitter balance and neurotrophic support, ultimately affecting the function and survival of Purkinje cells (*54*). Similarly, activated microglia can release pro-inflammatory cytokines, which can induce neuronal cell death pathways, including apoptosis and necrosis, when chronically elevated (*64*). Microglia has the ability to engulf and digest cellular debris, including dying neurons and their synapsis. Excessive removal of synapsis could also lead to synaptic dysfunction and ultimately contributing to Purkinje cells loss (*64*). This observation aligns with other neurodegenerative diseases characterized by neuroinflammation, such as multiple sclerosis, Alzheimer’s disease, and amyotrophic lateral sclerosis, where astrocytes and microglia are key regulators of the inflammatory response and diseases progression (*54*). We envision a possible scenario where partially impaired Purkinje cells may trigger Bergmann glia and microglia activation, which in turn could exacerbate Purkinje cell dysfunction, resulting in a vicious feed-forward cycle of Purkinje and granule cell neurodegeneration. Future studies will investigate how Sel1L deficiency in Purkinje cells leads to the activation of astrocytes and microglia in vivo.

Our findings indicate a key role of SEL1L-HRD1 ERAD in Purkinje cells in the development of cerebellar ataxia. This observation is consistent with a previous report in Finnish Hounds with cerebellar ataxia carrying a biallelic *SEL1L^S658P^* variant (*39*), which indeed leads to early-onset, non-progressive cerebellar ataxia in mice, as we recently demonstrated (*40*). Additionally, we have identified several patients with SEL1L M528R variant displaying ataxia, while others with SEL1L G585D or HRD1 P398L show mild hypotonia (*36*). The varying clinical presentations among these patients likely stem from differing degrees of ERAD dysfunction caused by these variants (*37*). Collectively, these studies underscore the potential pathogenic impact of SEL1L-HRD1 ERAD dysfunction. While none of these disease variants is linked to obvious UPR activation (*36, 37*), future studies are needed to delineate the underlying mechanisms, which may involve substrate- and cell type-specific effect contributing to this pathological process.

## MATERIALS AND METHODS

### Mice

Purkinje cell-specific *Sel1L*-deficient mice (*Sel1L^Pcp2Cre^*) were generated by crossing *Pcp2*-Cre mice on the C57BL/6J background (Jackson Laboratories #004146) with *Sel1L^fl/fl^* mice on the C57BL/6J background (*7*). Pcp-2 promotor begins within the first postnatal week (*48*) Age-and Gender-matched littermates were maintained in a temperature-controlled room on a 12-h light/dark cycle and used in all studies. All animal procedures were approved by the Institutional Animal Care and Use Committee of the University of Michigan Medical School (PRO00010658), and University of Virginia (4459), and in accordance with the National Institutes of Health (NIH) guidelines.

### Genotyping

Mice were routinely genotyped using PCR of genomic DNA samples obtained from ears with the following primer pairs: *Sel1L^fl/fl^*: F: 5’-CTGACTGAGGAAGGGTCTC-3’, R: 5’-GCTAAAAACATTACAAAGGGGCA-3’; Cre recombinase: F: 5’-ACCTGAAGATGTTCGCGATTATCT-3’, R: 5’-ACCGTCAGTACGTGAGATATCTT-3’.

### Behavioral studies

Most behavior procedures were performed by investigators blinded to the genotypes as previously described (*40*). For hindlimb clasping assessment, mice were lifted by tails and held over a cage for 1 min to assess abnormal hind limb clasping and scored as previously described (*49*). The footprints were analyzed for stride length (the distance covered by the same hind paw), stride width (the distance from one hind-limb that intersects perpendicularly with the line for stride length on the contralateral hind paw), paw matched distance (the distance between hind and forepaw), and the ratio between stride length and width. All mice received three trainings and a trial run. The balance beam study was used to evaluate motor coordination and balance as previously described (*40*). Mice were trained to stay upright and walk across an elevated narrow beam to a safe platform for two consecutive days (three times per day). On the third day, the time to cross 80 cm on the beam was measured. A video camera was set on a tripod to record the performance of the animals during the test. Room temperature, humidity, lighting, and background noise were kept consistent throughout the experiment.

### Western blot and antibodies

Tissues were harvested and snap-frozen in liquid nitrogen. The proteins were extracted by sonication in NP-40 lysis buffer (50 mM Tris-HCL at pH 7.5, 150 mM NaCl, 1% NP-40, 1 mM EDTA) with protease inhibitor (Sigma), DTT (Sigma, 1 mM) and phosphatase inhibitor cocktail (Sigma). Lysates were incubated on ice for 30 min and centrifuged at 16,000 g for 10 min. Supernatants were collected and analyzed for protein concentration using the Bio-Rad Protein Assay Dye (Bio-Rad). 20-50 mg of protein was denatured at 95°C for 5 min in 5x SDS sample buffer (250 mM Tris-HCl pH 6.8, 10% sodium dodecyl sulfate, 0.05% Bromophenol blue, 50% glycerol, and 1.44 M b-mercaptoethanol). Protein was separated on SDS-PAGE, followed by electrophoretic transfer to PVDF (Fisher Scientific) membrane. The blots were incubated in 2% BSA/Tris-buffered saline tween-20 (TBST) with primary antibodies overnight at 4°C: anti-HSP90 (Santa Cruz, sc-7947, 1:5,000), anti-SEL1L (home-made, 1:10,000) (*33*), anti-HRD1 (Proteintech, 13473-1, 1:2,000), anti-OS9 (Abcam, ab109510, 1:5,000), anti-IRE1α (Cell Signaling, 3294, 1:2,000), anti-PERK (Cell Signaling, 3192, 1:1,000), anti-Calbindin (Cell Signaling, 2173, 1:5,000), anti-GRP94 (Proteintech, 14700-1, 1:1,000), anti-BiP (Abcam, 21685, 1:5,000), GFAP (Cell Signaling, 3670, 1:2,000), anti-ATF4 (Cell Signaling, 11815, 1:1,000), anti-CHOP (Cell Signaling, 2895, 1:1,000), eIF2⍺ (Cell Signaling, 9722, 1:1,000), and p-eIF2⍺ (Cell Signaling, 9721, 1:1,000). Membranes were washed with TBST and incubated with HRP conjugated secondary antibodies (Bio-Rad, 1:10,000) at room temperature for 1h for ECL chemiluminescence detection system (Bio-Rad) development. Band intensity was determined using Image lab (Bio-Rad) software.

### RNA preparation and RT-PCR

Total RNA was extracted from tissues and cells using TRI Reagent and BCP phase separation reagent per supplier’s protocol (Molecular Research Center, TR 118). RT-PCR for *Xbp1* mRNA splicing was performed as previously described (*69*). The ratio of Xbp1s level to total Xbp1 (Xbp1u + Xbp1s) level was quantified by Image Lab (Bio-Rad) software. RT-PCR primer sequences are: mXbp1 F: ACGAGGTTCCAGAGGTGGAG, R: AAGAGGCAACAGTGTCAGAG; mL32 F: GAGCAACAAGAAAACCAAGCA, R: TGCACACAAGCCATCTACTCA.

### Histology

Anesthetized mice were perfused with 20 ml of 0.9% NaCl followed by 40 ml of 4% paraformaldehyde in 0.1 M PBS pH 7.4 for fixation. Brains were dissected and fixed overnight in 4% paraformaldehyde in PBS at 4°C. For hematoxylin and eosin (H&E) staining, samples were dehydrated, embedded in paraffin, and stained at the Rogel Cancer Center Tissue and Molecular Pathology Core or In-Vivo Animal Core at the University of Michigan Medical School. Quantification of Purkinje cell loss was performed on H&E-stained sections. Counts were normalized to the length of the Purkinje layer, as measured by Aperio ImageScope software, and shown as Purkinje cell density.

### Immunofluorescent staining

Paraffin-embedded brain sections were deparaffinized in xylene and rehydrated using a graded ethanol (100%, 90%, 70%, followed by a rinse in distilled water. Antigen retrieval was performed by boiling the slides in a microwave in sodium citrate buffer. Sections were then incubated in a blocking solution (5% donkey serum, 0.3% Triton X-100 in PBS) for 1 hour at room temperature and with primary antibodies anti-Calbindin (Cell Signaling, 2173, 1:100), anti-KDEL (Novus Biologicals, 97469, 1:200), anti-SEL1L (homemade, 1:100), anti-BiP (Abcam, 21685, 1:100), anti-GFAP (Cell Signaling, 3670, 1:100), and IBA1 (Novus Biologicals, NB100-1028, 1:100) overnight at 4°C in a humidifying chamber. The next day, following 3 washes with PBST (0.03% Triton X-100 in PBS), slides were incubated with the respective Alexa Fluor-conjugated to secondary antibodies (Jackson ImmunoResearch, 1:500) for 1 hour at room temperature, followed by mounting with VECTASHIELD mounting medium containing DAPI (Vector Laboratories, H-1500). Images were captured using a Nikon A1 confocal microscope at the University of Michigan Morphology and Image Analysis Core. Protein relative levels in immunofluorescent images were quantified using FIJI-ImageJ software (*65*).

### Transmission electron microscopy (TEM)

Mice were anesthetized and perfused with 3% glutaraldehyde, 3% formaldehyde in 0.1 M cacodylate buffer (Electron Microscopy Sciences, 16220, 15710, 11653). Cerebellum was dissected and cut into small pieces and fixed overnight at 4°C in 3% glutaraldehyde, 3% formaldehyde in 0.1 M Sorenson’s buffer (Electron Microscopy Sciences, 11682). The tissues were then prepared, embedded, and sectioned at the University of Michigan Histology and Imaging Core. Samples were stained with uranyl acetate/lead citrate and high-resolution images were acquired with JEOL 1400-plus electron microscope (JEOL).

### Statistical analysis

Statistics tests were performed in GraphPad Prism version 8.0 (GraphPad Software). Unless indicated otherwise, values are presented as mean ± standard error of the mean (SEM). All experiments have been repeated at least three times and/or performed with multiple independent biological samples from which representative data are shown. All datasets passed normality and equal variance tests. Statistical differences between the groups were compared using the unpaired two-tailed Student’s t-test for two groups or two-way ANOVA with Bonferroni’s multiple comparison test for multiple groups. *P* < 0.05 was considered statistically significant.

## AUTHOR CONTRIBUTIONS

M.T., H.W., and B.P. collaboratively designed and performed most experiments; L.L.L., H.H.W. and A.B.L assisted with some *in vitro* and *in vivo* experiments; Z.Z. provided insightful discussion; L.Q. directed the study and wrote the manuscript with help from M.T., H.W., and B.P.; B.P., H.W. A.B.L. and M.T. wrote the methods and figure legends; all authors commented on and approved the manuscript.

## ACKNOWLEDGEMENTS

We acknowledge members of the Qi and Arvan laboratories at the University of Michigan Medical School for technical assistance and insightful discussions; Dr. Andrew P. Lieberman (University of Michigan) for generously providing the Pcp2-Cre mouse; and the University of Michigan Animal Phenotyping Core for some of the behavioral tests (supported by P30 grants DK020572, DK089503 and 1U2CDK135066). This work was supported by RF1NS122060 (Z.Z.), 24AARG-D-NTF-1187603 and 1R35GM130292 (L.Q.). L.L.L. was supported in part by National Ataxia Foundation (NAF 918037).

## Data and materials availability

The materials and reagents used are either commercially available or available upon the request. All other data are available in the main text or Supplemental Materials.

